# Unraveling Biodiversity Change: A multi-scale decomposition of changes in European breeding bird diversity

**DOI:** 10.1101/2025.09.04.674078

**Authors:** M. Beck, P. Gauzere, F. Schrodt, W. Thuiller, IMPACTS consortium

**Author notes:** [authors in alphabetical order,]. first author (M. Beck).

## Abstract

**Aim:** Detecting and describing temporal changes in biological communities is fundamental to biodiversity research and applied ecology. Species richness remains a widespread metric in long-term monitoring, yet it obscures underlying processes since changes in species richness are often only the result of turnover, homogenisation and/or shifts in relative abundances. Furthermore, biodiversity trends and their drivers can vary across spatial scales, demanding spatially explicit approaches. This study aims at unravelling how changes in community structure shape trends in richness across spatial scales, offering a more mechanistic view on biodiversity trend detection.

**Location:** Europe

**Time period:** 1975 - 2023

**Major taxa studied:** Birds

**Methods:** We first assessed trends in breeding bird richness on local (site-level) and national scale for 25 European countries or sub-divisions using linear models. Next, we applied the multi-scale ‘Measures of Biodiversity’ (MoB) framework in a temporal context to decompose changes in species richness into contributions from individual density, species-abundance distribution, and con-specific spatial aggregation. We then quantify how these components drive species richness from local plots to national extents. Analyses were further conducted separately for farmland and forest guilds, as well as across ecoregions.

**Results:** Three general patterns emerged beyond variation among countries and functional guilds: Aggregation dominates local richness dynamics, evenness governs broad-scale trends, and density plays an intermediate role. Findings of distinct local and national trends in bird richness agree with previous findings, albeit we find more heterogeneous average trends among countries on local scales. Distinct trends and component’s patterns vary among ecoregions within countries, highlighting the need for sub-national analyses.

**Main conclusions:** This scale-explicit, component-based approach reveals how changes in community structure shape trends in species richness from local to national scales. Such mechanistic insights of biodiversity change might enable more precisely targeted conservation strategies and identification of external drivers.

## Introduction

Detecting and investigating temporal changes in biological communities is an important goal of biodiversity research and applied ecology (Magurran 2004; Moussy et al. 2021, Blowes et al. 2024; Dornelas et al., 2023). Assessing the total number of species (‘richness’) probably is the most straightforward metric and requires no complex data. Yet, declining long‐term losses in species richness (Newbold et al. 2015) or absence thereof on (Dornelas et al. 2014) are often symptoms of a range of underlying ecological processes, such as turnover, biotic homogenization or shifts in relative abundances. With univariate metrics they might go undetected. Thus, for a more in depth investigation of structural changes in communities, additional metrics complement the picture and address the multi-dimensionality of biodiversity change (Magurran 2004; McGill et al. 2007). They describe, for example, the relative distribution of abundances among species which can inform about dominance of few taxa (‘Pielous evenness’, ‘Species Abundance Distribution, SAD’), aggregation in space (‘clumping’, ‘patchiness’) or include other aspects of meta communities such as the regional species pool. Yet, truly understanding global change effects on biodiversity remains challenging.

A primary and often main obstacle to interpreting biodiversity trends is scale dependency: both diversity metrics and their environmental drivers vary across spatial scales (Chase et al., 2018; Chase et al. 2019). For example, the local richness of North American breeding birds has remained relatively stable in recent decades, whereas the richness across regions has increased; an apparently contradictory result when reported as such, but at second glance driven by species gains at broader scales (Chase et al., 2018). Similarly, Catano et al. (2021) demonstrated that the influence of the regional species pool on observed richness increases with increasing extent of sampling area. However, most empirical studies still focus on single scales - such as local plot(s), landscape polygon, or national inventory - which are often chosen arbitrarily or based on data availability. Lacking clear specifications of scale or unintentional comparisons of multiple scales can lead to misguided inferences about changes in biodiversity, ranging from over- or underestimation of trends or even contrasting conclusions about their direction (Catano et al., 2021; Janvier et al., 2022). Approaches developed in macroecology — including species–area relationships (SARs, e.g. Rosenzweig 1995) and scale-explicit models of species abundance distributions (SADs, e.g. McGill 2011) — provide conceptual and analytical frameworks to integrate spatial scale as a continuous variable in ecological analysis.

Second, despite the clear need for a multi-metric view on diversity (Santini et al. 2017) its implementation often remains limited. Empirical meta-analyses for example report covarying trends of richness and density (e.g. Sandal et al 2024, Davison et al. 2024, Rumschlag et al. 2023, Burner et al. 2021) or parallel declines in evenness and species loss (Zhang et al. 2021, Czarniecka‐Wiera et al. 2019). Others question the universality of these patterns (Soininen et al. 2012; Burner et al. 2021) and, overall, the interpretation often remains focussed on describing patterns rather than highlighting processes. The “Measurement of Biodiversity” (MoB) framework (McGlinn et al. 2019) addresses this aspect, by considering changes in species richness as a consequence of alternations in community structure in one or several aspects (addressed as ‘components’ by the authors and in the following) (He & Legendre 2002; McGlinn 2018). This allows partitioning species richness change into contributions from density (or total abundances), the shape of the SAD (including also regional richness), and spatial aggregation of individuals. The rationale is both ecological and probabilistic: All else being equal, sites with high densities or a more even SAD will also show higher species richness, simply by modulating the probability of encountering new species. For a similar reason, richness will be higher when individuals of each species are evenly aggregated in space, compared to highly clustered communities. When change in species richness per se is not the main interest, such a framework can reveal deeper changes in community diversity, driven by an interaction of the underlying community components. For example it identified SAD as the underlying component driving species richness along an elevational gradient (McGlinn et al. 2021) or under the impact of invasions (Kortz et al. 2021). However, it has not yet been applied in a temporal context despite its potential to reveal how diversity is shaped through time from local to large scales (Gauzere et al. 2025). Such temporal-spatial extension would shift the focus of trend detection from “How have richness, density, and evenness changed?” to “To what extent do changes in density, evenness, and aggregation explain observed trends in richness?”. Since community structure is likely to simultaneously change along in multiple components, estimating their relative contribution to changes in species richness will improve our understanding of diversity trends. For example, detecting increasing richness despite declining abundances (Rumschlag et al. 2023), could suggest that density was not the main driver of richness changes, or that the negative effect of abundance was compensated for by a positive effect of another, unmeasured, structural change.

Birds are widely monitored throughout the Northern Hemisphere, enabling extensive research on biodiversity trends and revealing highly heterogeneous patterns across local to continental scales (Leroy et al. 2023). The integration of species traits, functional guilds (Bowler et al. 2018, Rigal & Knape 2024, Gregory et al. 2023, Verhulst et al. 2004), or geographic context (Harrison et al. 2014) has advanced our understanding of population and community trajectories. While case studies have linked community-level changes to shifts in species richness, a general understanding of how such mechanisms operate across spatial scales—and whether universal patterns exist—remains elusive.

Here, we present the first study applying the MoB framework in a temporal context and across pan-European scales to decompose how changes in density, SAD-evenness and conspecific aggregation drive trends in breeding bird richness along continuous spatial scales. First, we quantify long‐term trends in bird richness and density at local (1 km^2^) and national scales, both overall and within functional guilds (farmland, forest), which are groups known to respond differently to human induced land use and cover change (Verhulst et al. 2004; Harrison et al. 2014; Rigal & Knape 2024). We ‘break’ arbitrary political boundaries to assess trends across ecoregions. Finally, we conduct a multi‐scale analysis that quantifies the contributions of density, SAD-evenness and aggregation to these changes in richness.

We expect that, considering the scale dependency of community metrics as well as driver effects, their effects on species richness temporal changes vary across scales. Due to its expected link to (local) habitat heterogeneity, we hypothesize that aggregation influences richness on small spatial scales. Density and SAD-evenness, subject to a large range of drivers ranging from local to large spatial scales, should be important at larger spatial scales. Farmland and forest guilds are expected to differ in the dominant components and/or the direction in which they influence richness, reflecting contrasting responses to land-use pressures and country-individual situations.

## Materials and Methods

### Data compilation

We assembled a European dataset of standardized breeding‐bird surveys coordinated under the Pan‐European Common Bird Monitoring Scheme (PECBMS). Participating national programs follow a common protocol, visiting each site once or twice per year during the core breeding season (April– June). The geographic extent of each survey generally is the entire country area, with the exceptions of Belgium and Catalonia, where sub-divisions are covered. Sampling site selection varies among countries, with some applying a stratified or grid-based random sampling design with assigned observers, while others rely on more opportunistic approaches where observers choose their own survey locations. For each site, subplots are visited, which are placed to cover the full range of local habitats either within a fixed radius of 1-4km^2^ (‘point counts’, 15 surveys) or along a defined transect of 1-4.5km (mean 2.7km, 8 surveys). In schemes following ‘territory mapping’ (Netherlands, Switzerland) selection of sites and subplots within was more systematic, attempting to cover the whole country area (Swiss Breeding Bird Atlas 2013–2016). At each subplot, observers record all individuals seen and/or heard at species level. For each site and year, subplot counts were summed and the maximum annual count per species across visits was retained. For a few surveys, this aggregation was based on the mean (n=1, Slovakia) or varies among species (but maximum for most species) (n=3, Austria, Netherlands, Bulgaria see Tab. S1 for details). In the following, the term ‘abundance’ refers to these standardized counts, acknowledging that BBS data do not correct for detectability. Records outside of the published species breeding-range maps (European Bird Atlas I & II; Hagemeijer & Blair, 1997; Keller et al., 2020) or flagged by national coordinators for confidentiality reasons (e.g. rare or protected species) were excluded. Despite the same general sampling procedure, we analysed the national surveys separately to avoid obscuring differences due to e.g. different densities in sampling sites, varying temporal scales or country-specific differences in sampling realisation.

### Site selection and data filtering

To ensure reliable trend estimates, we only retained sites with at least five years of sampling data. We removed rare species recorded in fewer than 0.1% of site-years or accounting for fewer than 0.01% of all individuals, and excluded site-years with fewer than five total individuals to avoid distortions caused by very small samples. Annual abundances above the 95^th^ percentile across total abundances were winsorised i.e. set to the value of that threshold to reduce the influence of extreme flock events. As sampling intensity varied among years and schemes, we rarefied each survey’s sites to equalise the number of sites per year. Bootstrapping was implemented around this site selection (n = 100) and final outcomes of the analyses were averaged over iterations with confidence intervals reflecting the variability.

Our final dataset consisted of 25 surveys across 23 countries (two schemes in Belgium and Spain) spanning 1975–2023 (mean time period of 24.9± 11 years). Each survey comprised 19–5843 single sites (mean = 657± 1171; median = 267) that were observed 5-49 times (mean=13.3± 7.1; median=12). In total 384 species were recorded (mean = 132± 25 species per survey, median=137) and species occurred on average in 8.6± 8 surveys (median=5; 30% of species were observed in half of the surveys). The location of each site was defined by its geographic centroids in our spatial analyses. More information including a map of the sites can be found in the supplementary material (tab. S1, Fig. S2).

### Functional guilds and ecoregional stratification

To test whether community‐structural drivers differ among ecological guilds, we classified each species as farmland or forest, based on PECBMS methods (PECBMS 2024). All analyses were run on the full community and separately by guild.

We also examined within‐country variation among ecoregions (Metzger et al. 2005), retaining only those regions allowing to analyse ≥ 12 sites per year (the minimum in the country‐level analyses). To reduce within-ecoregion spatial discontinuities and to increase sample size, we merged the “Mediterranean North” and “Mediterranean South” zones into a single “Mediterranean” region (Fig. S2).

### Trend estimation at α- and γ-scales

Following our focus on scale dependency, we calculated α richness (number of species per site) and γ richness (total number of species per country or ecoregion) for each year. As Davison et al. 2024 already highlighted that bird communities showed linear trends on local and global scale, we applied a linear trend approach to detect changes in the diversity time series: For alpha level, we fitted linear mixed-effects models with ‘richness’ as the response, ‘year’ as a fixed effect, and ‘site’ as a random effect to account for confounding site characteristics. On gamma scale, we fitted simple linear regressions of annual richness against year, since only one value is available per year.

### Decomposing richness change using MoB

Following MoB framework, the contribution of density (‘N’), species-abundance-distribution (‘SAD-evenness’) and conspecific aggregation (‘Agg’) to the observed changes in species richness across space were quantified using mobr R package (McGlinn et al. 2018) in R version 4.4.1 (ref).

The approach uses three types of rarefaction obtained by randomly shuffling samples and/or (georeferenced) sites that describe species richness across sampling effort and contain information about one to all three components. For example, a random sampling of plots ignoring their spatial distribution will contain information about density and evenness, whereas a sampling of sites keeping their spatial distribution (i.e aggregating neighbours sites first) will additionally contain information about individuals’ spatial aggregation. For each type of sampling, there is one rarefaction curve per year, so that changes over year can be calculated as the difference between the curves. To isolate the effects of N, SAD-evenness and Agg, the three types of curves are subtracted from one another for the same year, resulting in one final rarefaction curve for each component. Analysing the change through time (i.e. difference between individual curves within one type) at a defined subset across sampling effort allows to map the change in richness over time at each spatial scale due to the respective component. The linear slopes of these curves then quantify the contribution of N, SAD-evenness and Agg over time on species richness. For more details we refer to the official documentation of the framework (McGlinn et al. 2019).

Practically, this was done by applying the function *get_delta_stats()* in the mobr package treating ‘year’ as the environmental variable that defines a gradient along the sites (in this case temporal gradient). The efforts (i.e. the spatial scale) at which effects are estimated were automatically chosen equally spaced on logarithmic scale. N=1000 permutations were run to obtain a Null model for each of the three rarefaction curves.

To facilitate cross-country comparisons, the component effects on species richness were summarised across a range of spatial scale continuum to represent local (sampling effort < 5% quantile of maximum sampling effort), intermediate (40-60% quantile) and large (> 95% quantile) spatial scales. Only significant effects values passing the Null model were considered in mean calculation.

All scripts are available on GitHub (), bird data is available upon request from PECBMS or national coordinators.

## Results

### Species richness at local and national scales

Across the 25 European surveys, temporal trends in local (α-scale) taxonomic richness were highly variable (Fig. 1; Table S3). Fifteen surveys exhibited significant increases in mean site-level richness, while ten showed declines. The steepest gains occurred in Latvia (+0.28 species yr^−1^), and the largest losses in Croatia (−0.40 species yr^−1^). On average, local richness rose modestly by +0.03 species yr^−1^. Trends within forest birds mirrored the overall pattern (16 surveys increasing, eight decreasing; Fig. S2), but farmland birds displayed predominantly negative local trends (21 of 25 surveys declining) except for Cyprus, Estonia and Switzerland where local richness increased. Farmland bird richness changed on average by −0.04 species yr^−1^ (+0.04 species yr^−1^ in Cyprus to -0.23 species yr^−1^ in Slovakia); forest bird richness +0.02 species yr^−1^ (+0.11 species yr^−1^ in Latvia to -0.06 species yr^−1^ in Slovakia). At the national (γ) scale, trends in richness were almost uniformly positive (Fig. 1; Table S3). 20 surveys showed significant increases (mean +0.29 species yr^−1^), with the strongest occurring in Estonia (+0.81 species yr^−1^). Only in Bulgaria (−0.74 species yr^−1^) and Switzerland (−0.03 species yr^−1^) richness declined. On average richness increased by +0.28 species yr^−1^.

**Fig. 1:**
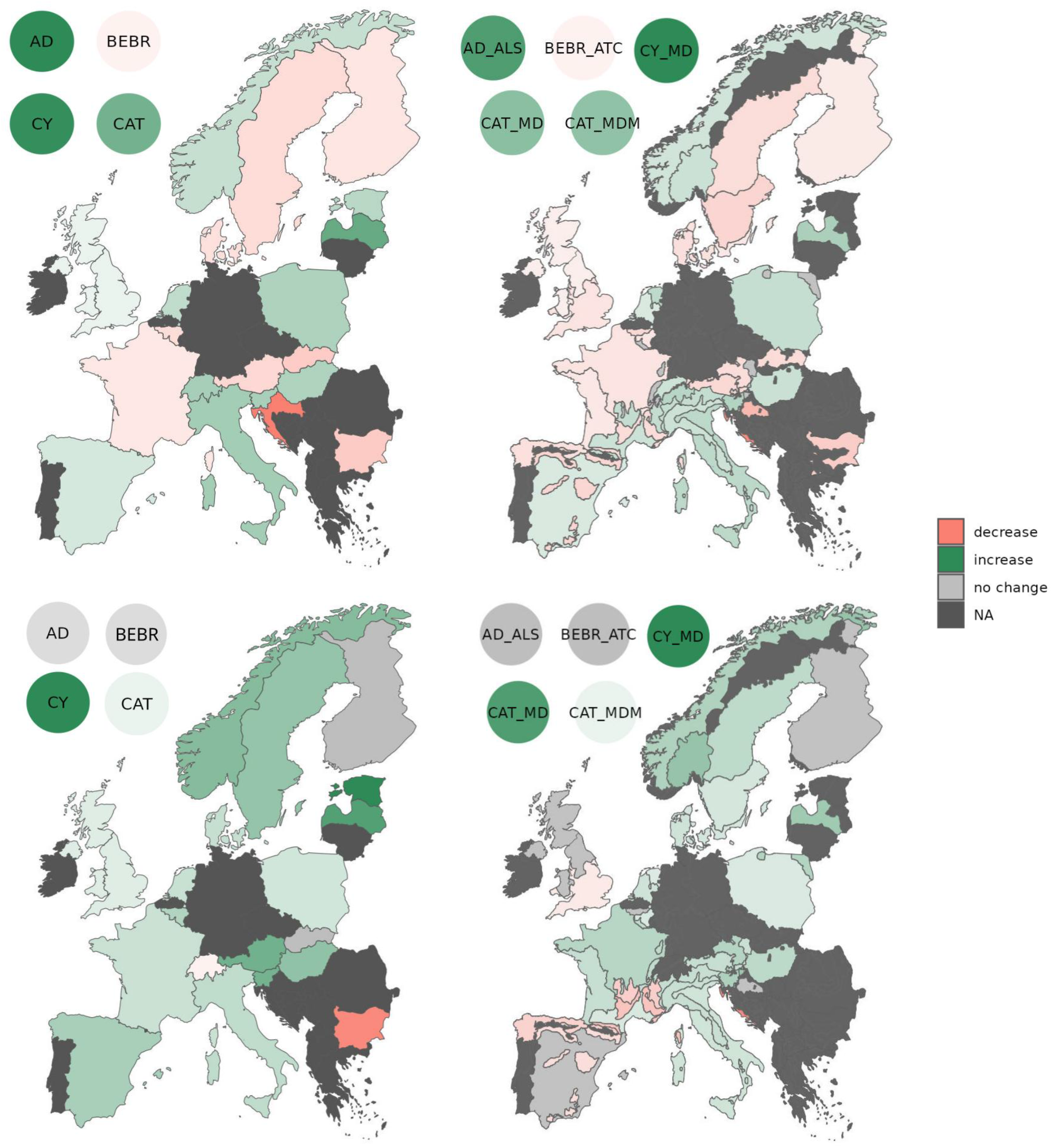
Direction of significant trends in taxonomic richness at local (top) and national (bottom) scale for the overall community on a national scale (left) and ecoregions therein (right) as obtained from linear mixed models accounting for site as random factor or simple linear model. For visibility the results of small-area surveys are shown in separate circles (AD=Andorra, BEBR=Belgium-Brussels, CY=Cyprus, CAT=Spain-Catalonia;). Color indicates decrease (red), increase (green) or no change (gray); color intensity corresponds to the slope of change. Dark gray areas were not included in the study. significance at alpha 0.05

The picture for the functional guilds was more heterogeneous. Farmland bird richness increased in eleven and declined in nine surveys with strongest trends occurring in Cyprus (+0.17 species yr^−1^) and Hungary (-0.23 species yr^−1^). Trends in forest birds were positive in ten and negative in four surveys, ranging from +0.22 species yr^−1^ in Hungary to -0.22 species yr^−1^ in Slovakia. Average changes were +0.01 and +0.03 species yr^−1^ for farmland and forest guilds.

For both total and guild richness national trends were stronger than local trends. On both spatial scales, changes detected in a given survey in functional guilds were weaker than for the complete community for most surveys.

Within-country ecoregions were largely uniform and reflected national trends (Fig. 1). The only exceptions are France and Spain, where bird richness in the Mediterranean and/or Mediterranean mountains showed opposing trends to the rest of the country on both spatial scales.

### Component effects on changes in richness

At small spatial scales, trends in richness were mostly driven by changes in aggregation and density in 12 and 11 surveys, respectively (Fig. 2, Fig. 3, Tab S4). On average and irrespective of its direction, aggregation lead to changes of 0.10 ± 0.10 species yr^−1^, ranging from +0.15 (Bulgaria) to -0.44 species yr^−1^ (Austria). The average strength of density effect was 0.06 ± 0.06 species yr^−1^ (+0.11 species yr^−1^ in Cyprus to -0.29 species yr^−1^ in Hungary). Changes in SAD-evenness played a smaller role (mean +0.03 ± 0.04 species yr^−1^). For all metrics the direction of their temporal change and thus their effect on richness was heterogeneous among surveys, but positive effects - resulting from increased density, more even SAD or more evenly distributed individuals - predominated, especially for SAD (n =17). At mid scales, changes in SAD-evenness (12 surveys) or aggregation (14 surveys) accounted for most change in richness with the average strength of SAD-evenness effect (0.27 ± 0.16 species yr^−1^, ranging from +0.54 in Estonia to -0.47 in Bulgaria) was stronger than that of aggregation (0.10 ±0.07 species yr^−1^, ranging from +0.26 in Andorra to -0.19 in Italy). Average density effect accounted for changes of 0.08 ± 0.07 species yr^−1^, but it was only significant in 7 surveys (strongest positive and negative effects in Cyprus (+0.19) and Norway (-0.12)). SAD-evenness effect on richness was predominantly positive across surveys (suggesting more even SADs), while aggregation effect was negative (suggesting increased levels of aggregation).

**Fig. 2:**
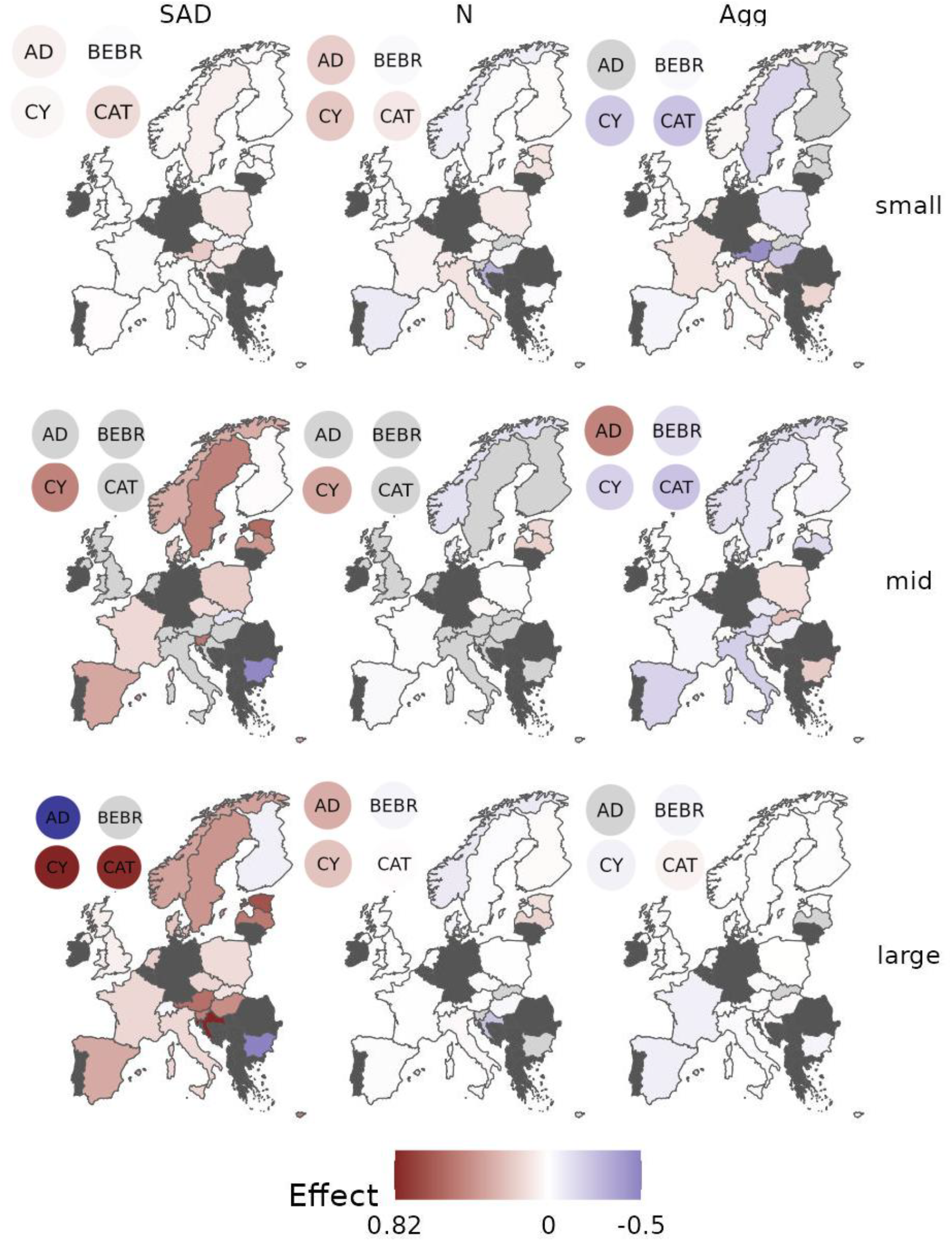
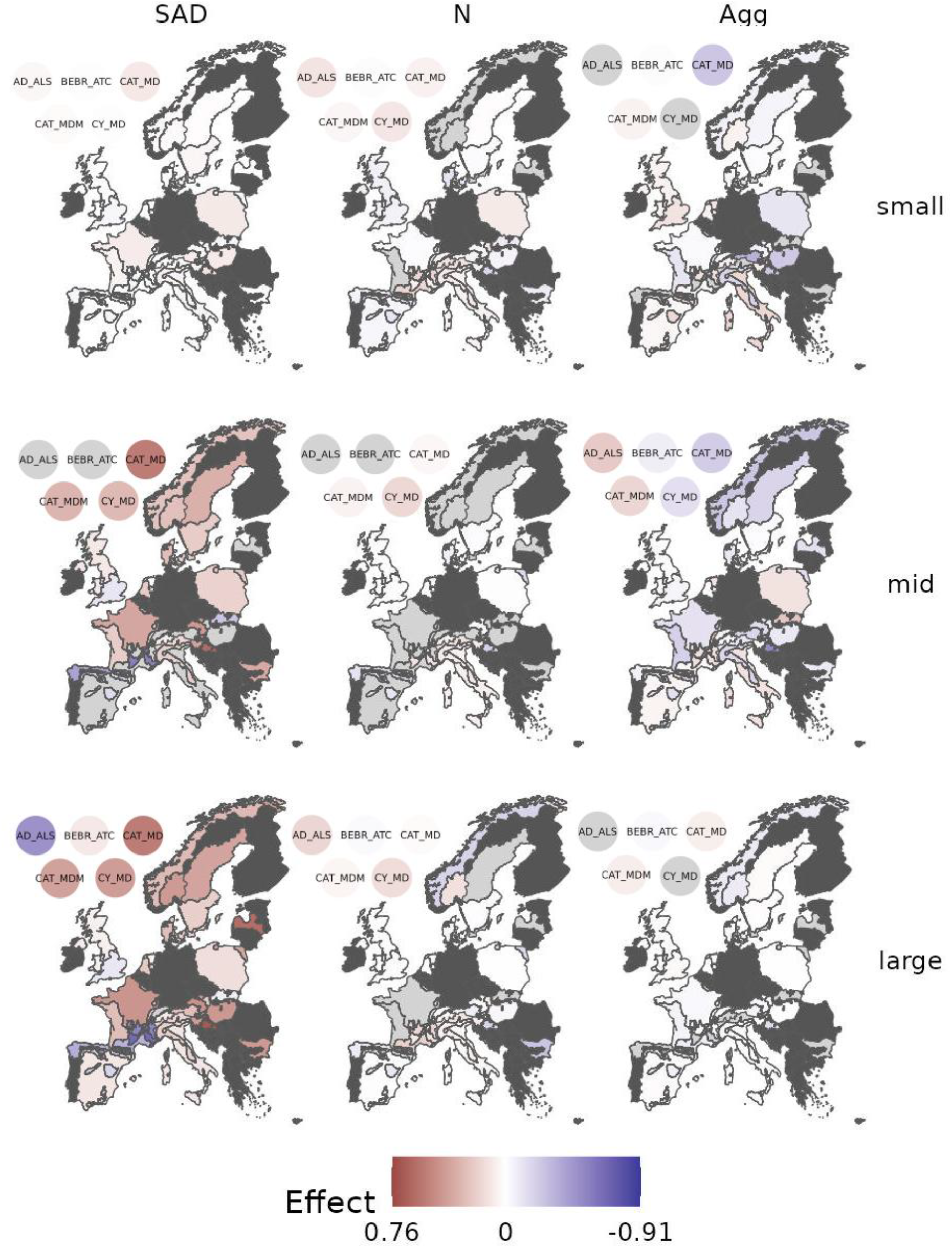
Mean effect of SAD-evenness, N and aggregation on species richness of the overall community at small, mid and large spatial scales for each sampling scheme (top) and individual ecoregions therein (bottom). Color indicates the direction and strength of the effect. Gray colored areas indicate the absence of significant effect (effect size not passing Null model), dark gray colored areas were not included. For visibility the result for small surveys is shown in labelled circles (AD=Andorra, BEBR=Belgium-Brussels, CY=Cyprus, CAT=Spain-Catalonia).

**Fig. 3:**
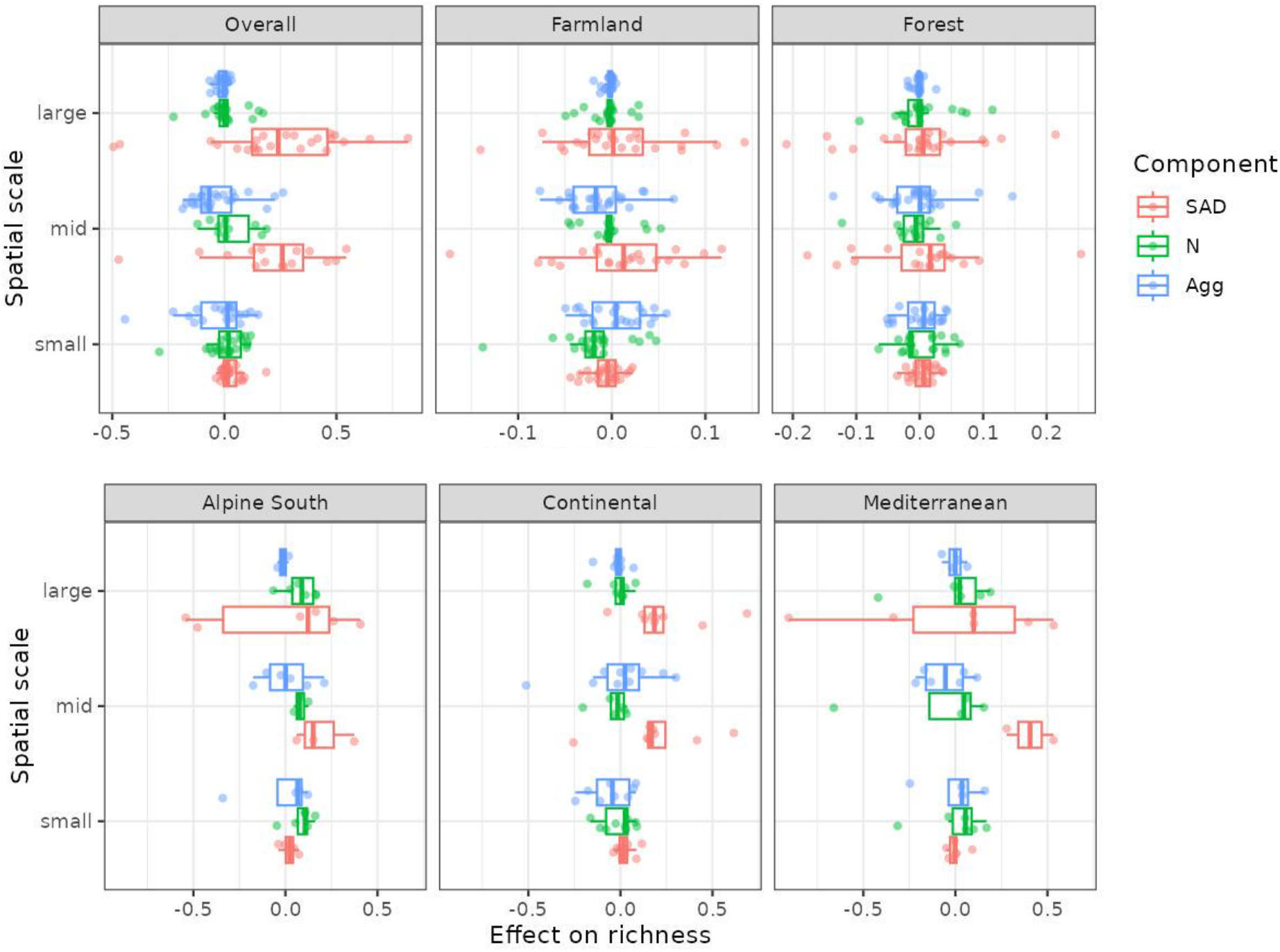
Mean effects of SAD-evenness, N and aggregation (‘Agg’) on species richness at small, mid and large spatial scales for the overall community and functional guilds (top) as well as the overall community in most frequent within-country ecoregions (bottom). Boxplots show median and quantiles (25th, 75th), points show raw mean effects. Please note that individual scaling of x-axis to facilitate visibility. Additional with-country ecoregions are provided in the S5.

On large spatial extents, SAD-evenness drove richness trends in 24 of 25 surveys, accounting for an average change of 0.33 ± 0.21 species yr^−1^, but also showing the largest variation (Fig. 3). Effect strength ranged from +0.82 species yr^−1^ in Croatia to -0.49 species yr^−1^ in Bulgaria. Density and aggregation on average accounted for changes of 0.05 ± 0.06 and 0.02 ± 0.02 species yr^−1^, respectively. While SAD-evenness effects were positive in most surveys (n = 20), the direction of density and aggregation effects were more heterogeneous among surveys. It is to be noted that at largest spatial scales the aggregation effect is supposed to approach zero since all sites are accumulated (McGlinn et al. 2019).

In the functional guilds the relative importance of component effects across spatial scales, overall, was similar to those on the overall community (Tab. S5, Fig. S6). Some noteworthy differences include farmland birds, whose richness on small scales was predominantly negatively impacted by changes in density (declines) (17 surveys). Further, while change in SAD-evenness was the strongest driver of species richness on large scales also for functional guilds, its effect was more heterogeneous and caused richness declines in some surveys. Overall, effects were weaker in both functional guilds than for the overall community.

Within-country ecoregions showed similar directions of component effects in most surveys, but the strength of these effects often varied. In a few cases—such as in France, Spain, and Italy—shifts in SAD-evenness or aggregation increased species richness in one ecoregion while contributing to declines in another. In other cases, significant and meaningful effects of a given component on richness were restricted to a single ecoregion. Across surveys, the component effects within the same ecoregion varied widely, both in magnitude and direction (Fig. 3; Fig. S5). Compared to national analyses, more regional cases showed non-significant results; however, the strength of detected effects was generally within the same range as those observed at the national level.

Raw component effects across continuous spatial scales are provided in S8.

## Discussion

Our work reveals the underlying drivers of temporal trends in bird richness across scales and demonstrates how this can add a more mechanistic view on biodiversity trend detection. The spatial scales (small-mid-large) addressed in the multi-scale part are to be seen relative to the countries’ area and the sampling site coverage. Nevertheless, as a scale addressed by monitoring or management programs it remains useful. Similarly, the study period differs between surveys and thus may address slightly different change trajectories.

Bird richness significantly changed in all countries or sub-divisions considered in this study, both locally and on a national scale. The direction of change - increase, decline - is very heterogeneous among surveys on local scales, while on national scales overall bird richness increased in almost all countries. A systematic review already revealed a predominance of positive trends in bird richness on local and regional scales (Leroy et al. 2023) despite also a large part of studies reporting stable average species numbers, yet masking large variations among individuals sites (e.g. Dornelas et al. 2014). Ecologically, a likely explanation stems from the fact that, especially on larger spatial scales, extinctions happen slower than species gains due to arrival of new taxa (see e.g. Leroy et al 2023).

The findings differ for functional guilds. Local richness of farmland birds was in decline in most European countries, which is in line with trends in total abundances (see SM) and previously reported population and community trends throughout Europe (Rigal 2023, Voříšek et al. 2008, Burns et al. 2021). While previous population analyses suggest mixed (Rigal 2023, Voříšek et al. 2008) or no/lower (Burns et al. 2021) trends for European forest birds, we find their richness increasing in most countries. The absence of significant trends on national scale suggest a rather stable species inventory of forest and farmland birds in some countries albeit changes happening on local scales. The second part of our analysis will explore which structural changes in the communities drove these trends.

Changes in bird richness were associated with shifts in all three components - evenness of the species pool, the density of individuals and their conspecific spatial aggregation - with the dominant driver varying with spatial scale. The effects of all three components were strongly scale-dependent and varied among surveys, yet three patterns emerged across most surveys and the functional guilds: (1) Changes in density rarely drove species richness except at small or mid spatial scales (sometimes) (2) The aggregation effect was variable across spatial scales, but governed many species losses or gains at local to mid range. (3) Changing SAD-evenness had a consistent positive effect at the national level.

The importance of the density (sampling) effect on species richness, or ‘more individuals hypothesis’, has already been pointed out especially for smaller spatial scales for a range of taxa (e.g. Stoch et al. 2018, Palmer et al. 2008, Feng et al. 2022, Morin et al. 2025), thus matching our finding. The simple decline (or increase) of local abundances often was the main driver of local diversity losses (or gains).

Aggregation can be linked to configuration of the habitat (Green et al. 2007, Plotkin et al. 2000) and it has already been identified as a main driver of bird richness on small or intermediate spatial scales (Būhning-Gaese 2007). A very heterogenous, local site might harbor high biodiversity by offering various niches. However, in a multi-site perspective as taken by our approach, sites with distinct configurations leading to clustering of species among these sites, actually reduce their detection probability and expected species richness in a random sample. In a homogenous landscape, individuals of a given species are likely more evenly spread across communities of neighboring sites, thus increasing their detection probability. Highly variable habitat configuration (and changes thereof) within and among European countries likely explains the variable changes in aggregation patterns with both positive and negative consequences for species richness we observed. Aggregation can also be determined by species identity. Behaviours such as flocking or cooperative breeding can favor aggregation (Bourque & Desrochers 2006, Salek et al. 2022, Gotelli et al. 2010), while intra-specific territoriality or competition can reduce it (Gotelli et al. 2010). However, those traits are very species-specific suggesting habitat configuration as the main source for varying aggregation. For other taxa, notably plants, poor dispersal abilities causing aggregation (Green et al. 2007) might be more important.

With increasing spatial scale shifting SAD-evenness, i.e. the equilibrium between dominant and rare species as well as the regional species pool, gains importance, reflecting selective population responses and regional pool dynamics. Throughout Europe, increasing evenness contributes to overall species gain at large scales, indicating coexistence of rarer taxa on national scales (Hillebrand et al. 2008) or declines in common species (Burns et al. 2021, Inger et al. 2014). In contrast, local dominance was rather stable and thus did not contribute much to re-structuring bird communities. In (some) Mediterranean ecoregions and within functional guilds, notably farmland birds, however, declining evenness negatively impacted bird richness across all scales. Other works applying MoB already identified SA-evennessD as the most important driver across all spatial scales for fish richness in protected vs. unprotected sites (Blowes et al. 2019), plant richness after pine invasion (Kortz et al. 2021) or tree diversity among regions (Feng et al. 2022). We showed that throughout Europe it is also the main driver of bird richness over time on large scales.

Trends in species richness and the associated effects of SAD-evenness, density and aggregation, were weaker for farmland and forest guilds than for the overall community. This likely is due in part to the smaller species pool within the guilds, but may also reflect the historical shaping of community structure (Le Provost et al. 2020 (?)) resulting in relatively stable farmland forest species pool in recent years. The latter however contrasts with reports suggesting that habitat specialists are more vulnerable to declining diversity (Morelli et al. 2020, Gregory et al. 2007).

The component-based approach adds to classical richness change detection in two important ways: 1) unraveling the interplay of multiple effects and 2) facilitating the identification of external (environmental, anthropogenic) drivers.

Understanding how shifts in SAD-evenness, density and aggregation each shape species richness offers a more mechanistic understanding of their possibly concurrent influence: In some surveys (e.g. Cyprus, Austria, Spain-Catalonia), negative effects of increasing aggregation were counteracted by increasing abundances resulting in only small net changes in richness. Decreasing aggregation and increasing abundances as on local scales in France and Italy suggest a gain in species richness due to increases in common rather than specialized birds (assuming specialist birds to show higher aggregation patterns). The inverse was observed in Spain, where increased aggregation and reduced densities can suggest a loss mainly in common species (already observed by Inger et al. 2019). However, we can also imagine an overall loss across all species under a strong external driver forcing higher aggregation (e.g. extreme habitat modification), highlighting the need for detailed understanding of environmental pressures, but also how such component-decomposition can direct further analysis.

Identifying the community-level processes underlying changes in species richness enables a more targeted assessment of the external forces driving these trends. This is essential for robust attribution studies and effective conservation planning. For example, if local richness declines are primarily driven by changes in spatial aggregation, conservation efforts can focus on external pressures that affect this component, such as habitat homogenisation resulting from land-use change. If we want to attribute the drivers of species richness at broader spatial scales, we can explicitly focus on drivers acting on SAD-evenness (e.g. climate-induced range shifts or land-cover change disproportionately affecting common species).

Geographic heterogeneity of trends in bird richness as well as of the relative importance of community-structuring mechanisms could not be explained by data or sampling characteristics (SUPP/not shown). Also, the absence of clear latitudinal and/or longitudinal patterns suggests region-specific drivers beyond the climate gradient or historical land-use (Donald et al. 2006, Verhulst et al. 2004) and a complex relationship between the environment and the way community metrics govern species richness. Opposing trends in component effects between within-country ecoregions (e.g. in France, Spain, Italy) and species richness trends at the national scale that seem to be driven by single ecoregions (Spain, France) highlight that even a simple secondary layer on sub-national scale reduces large-scale landscape heterogeneity and can drastically change the detection and interpretation of trends (Harrison et al. 2014)

This framework is easily adaptable to other facets of diversity (e.g. functional, phylogenetic, genetic) where corresponding data is available. Broad application of this approach is mainly limited by its requirement for highly structured community abundance data. This data unfortunately is often not available outside commonly monitored taxa which highlights the need to share raw data.

## Conclusion

Our analysis revealed how changes in SAD-evenness, density and spatial aggregation shape species richness through time from local to national scales. Changes in density or aggregation are main drivers of bird richness on small or intermediate spatial scales, whereas the effect SAD-evenness clearly dominates at large spatial scales. Despite heterogeneity among European countries, ecoregions and functional guilds three common patterns emerge, suggesting some generality of these findings. In addition, our study highlights the importance of spatial aggregation for community structure which, until now, gains less attention with regard to diversity change. Such mechanistic insights into community structuring through time facilitate the identification of external drivers causing biodiversity trends, which is the basis for robust attribution or targeted conservation actions.

## Acknowledgements

Our biggest thanks go to all volunteers who, from year to year, participated in the collection of bird count data. We also thank the national organisations for collecting and maintaining the data, and above all sharing it. […] We further thank CESAB for welcoming and hosting our workshops.

## Code and data availability

All code used to run the analysis and visualise the results is available on GitHub (github.com/Sikaiana/MacroDyn). Bird data is available upon request from PECBMS or national coordinators.

