## Supplementary material for "Unraveling Biodiversity Change: A multi-scale decomposition of changes in European breeding bird diversity"

*S1: Characteristics of surveys. Year indications refer to the overall survey and do not provide details about individual sites. Sampling indicates type of transect (point, line) or territory mapping.*

| Dataset | cd | Year_ start | Year_ end | Year_ period | nsite/ year | mean_ts length | Sites_density [nsites/year/100 km2] | Sampling |
|---|---|---|---|---|---|---|---|---|
| Andorra | AD | 2011 | 2021 | 11 | 14 | 9.37 ± 2.19 | 29.91453 | line |
| Austria | AT | 1998 | 2023 | 26 | 142 | 12.11 ±6.4 | 1.69308 | point |
| Belgium-Brussels | BEBR | 1992 | 2022 | 31 | 54 | 24.1 ± 6.2 | 335.40373 | point |
| Belgium-Wallonia | BEWA | 1990 | 2022 | 33 | 40 | 12.24 ± 6.8 | 2.37473 | point |
| Bulgaria | BG | 2005 | 2021 | 17 | 36 | 8.21 ± 2.9 | 0.32449 | line |
| Cypres | CY | 2006 | 2023 | 18 | 22 | 10 ± 3.3 | 2.37812 | line |
| Czechia | CZ | 1982 | 2021 | 40 | 44 | 12.38 ± 7.11 | 0.55788 | point |
| Denmark | DK | 1976 | 2023 | 48 | 34 | 12.11 ± 7.6 | 0.79215 | point |
| Estonia | EE | 1983 | 2021 | 39 | 20 | 10.71 ± 6.3 | 0.44116 | point |
| Spain-Catalonia | CAT | 2002 | 2023 | 22 | 139 | 13.47 ± 5.1 | 4.33143 | line |
| Spain | ES | 1998 | 2022 | 25 | 191 | 10.31 ± 5.1 | 0.40305 | point |
| Finland | FI | 1984 | 2022 | 39 | 32 | 13.46 ± 9.1 | 9.45E-05 | point |
| France | FRA | 2002 | 2019 | 18 | 551 | 9.68 ± 3.6 | 1.01298 | point |
| Croatia | HR | 2014 | 2021 | 8 | 51 | 7.38 ± 0.6 | 0.90116 | point |
| Hungary | HU | 1999 | 2022 | 24 | 108 | 9.8 ± 5 | 1.16084 | point |
| Italy | IT | 2000 | 2023 | 24 | 124 | 12.8 ± 5 | 0.4115 | point |
| Latvia | LV | 2005 | 2021 | 17 | 12 | 9.5 ± 3.6 | 0.18578 | line |
| Netherlands | NL | 1984 | 2023 | 40 | 231 | 14.08 ± 9.7 | 5.5605 | territory |
| Norway | NO | 2007 | 2022 | 16 | 97 | 10.49 ± 3.1 | 0.29921 | point |
| Poland | PL | 2007 | 2023 | 17 | 479 | 11.92 ± 3.8 | 1.53184 | line |
| Slovenia | SI | 2008 | 2023 | 16 | 72 | 11.31 ± 3.4 | 3.55152 | line |
| Slovakia | SK | 2005 | 2021 | 17 | 18 | 9.08 ± 4.2 | 0.36708 | point |
| Sweden | SWE | 1975 | 2023 | 49 | 44 | 13.98 ± 8.6 | 9.83E-05 | point |
| Switzerland | SWISS | 2001 | 2023 | 23 | 261 | 22.8 ± 0.5 | 6.32099 | territory |
| United Kingdom | UK | 1994 | 2023 | 30 | 632 | 15.12 ± 7.1 | 2.59431 | line |

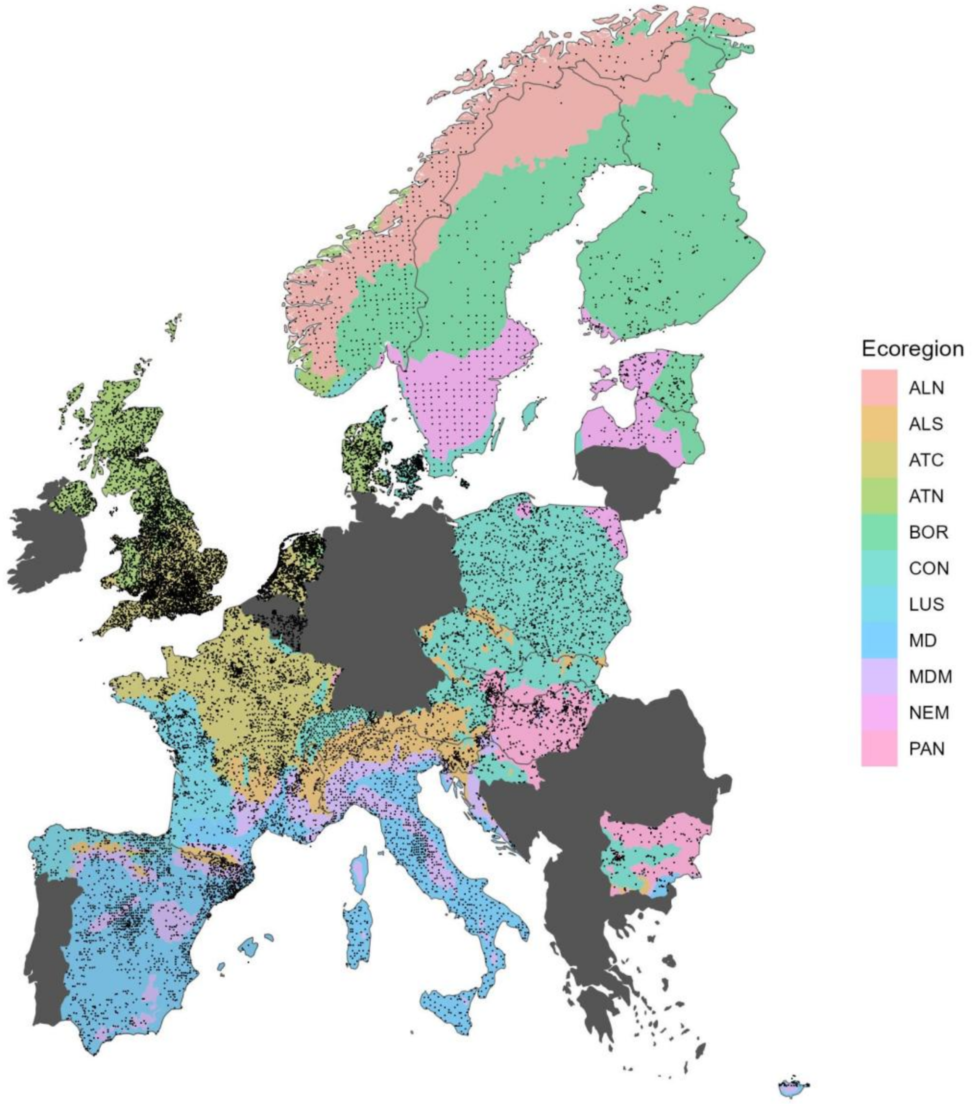


*S2: Map of final dataset. Each black point represents one sampling site; colors code ecoregions as defined by Metzger et al. (2005): Alpine North (ALN), Alpine South (ALS), Atlantic Central (ATC), Atlantic North (ATN), Boreal (BOR), Continental (CON), Lusitanian (LUS), Mediterranean (MD), Mediterranean Mountain (MDM), Nemoral (NEM), Pannonian (PAN).*

*S3: Direction of significant trends in taxonomic richness at local (top) and national (bottom) scale for the farmland (left) and forest (right) guild as obtained from linear mixed models accounting for site as random factor (local scale) or simple linear model (national scale). Color indicates decrease (red), increase (green) or no change (gray); color intensity corresponds to the slope of change. Dark gray areas were not included in the study.  significance at alpha 0.05*

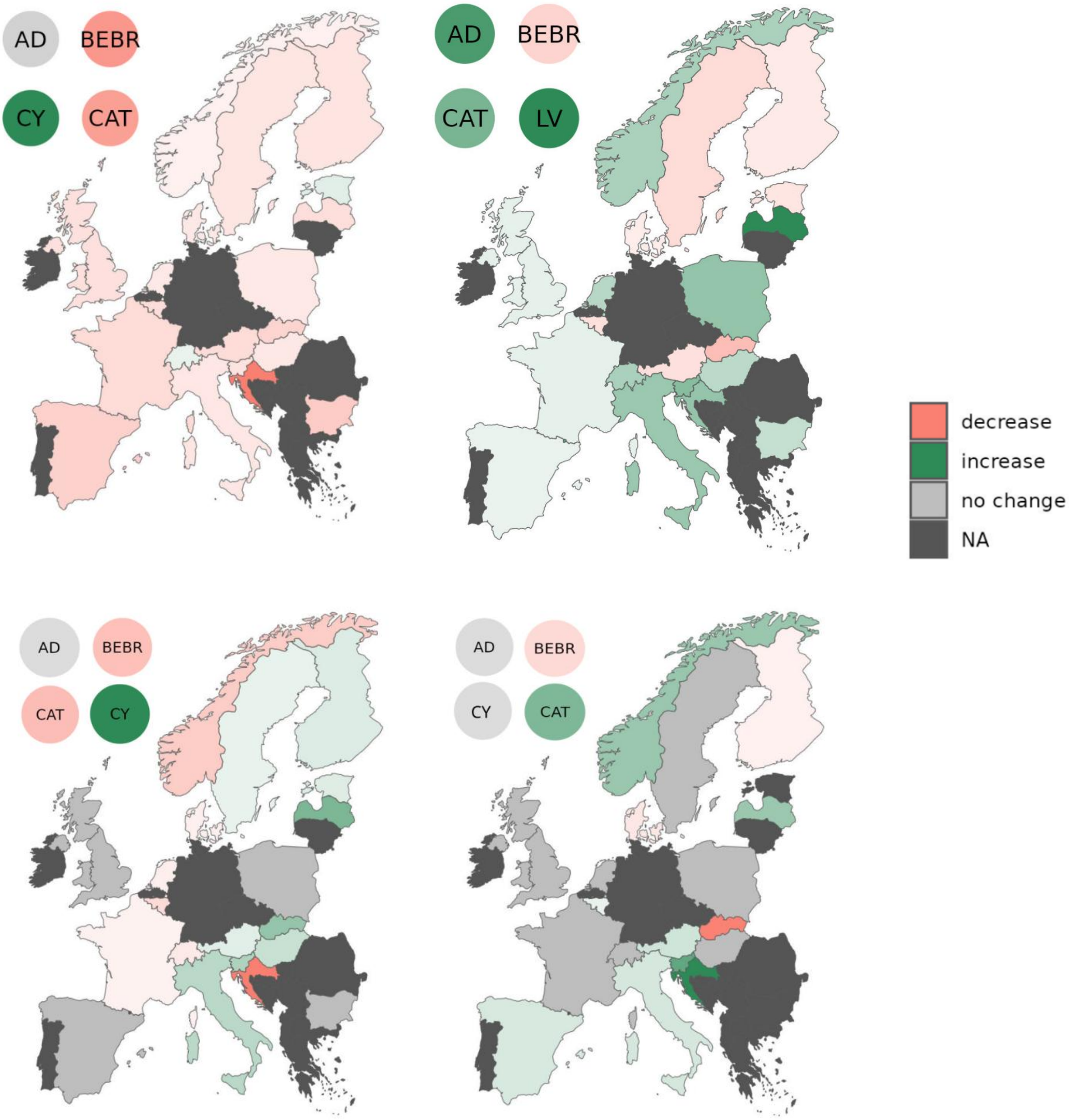

*S4 Mean slopes of trends in richness on alpha and gamma scale for each survey and functional guild. Only significant slopes from bootstrap runs have been included in calculation of mean; absence of values indicates that in none of individual bs trend was significant*

| | overall richness | | farmland richness | | forest richness | |
|---|---|---|---|---|---|---|
| country | α | γ | α | γ | α | γ |
| AD | 0.2131 | NA | -0.0558 | NA | 0.0841 | NA |
| AT | -0.0956 | 0.5277 | -0.051 | 0.0237 | -0.0267 | 0.0473 |
| BEBR | -0.0318 | NA | -0.018 | -0.0574 | -0.0122 | -0.1221 |
| BEWA | -0.0807 | 0.2634 | -0.0493 | -0.0574 | -0.0261 | 0.0195 |
| BG | -0.1399 | -0.7403 | -0.0841 | -0.1397 | 0.0311 | NA |
| CAT | 0.1221 | 0.4879 | -0.0369 | -0.044 | 0.0431 | 0.0331 |
| CZ | 0.0729 | 0.1569 | -0.0203 | 0.0271 | 0.0323 | 0.0098 |
| CY | 0.2034 | 0.6102 | 0.0392 | 0.1723 | NA | NA |
| DK | -0.0675 | 0.1815 | -0.0275 | -0.0191 | -0.0169 | -0.0382 |
| EE | 0.1095 | 0.8072 | 0.0177 | 0.0276 | -0.0226 | NA |
| ES | 0.0495 | 0.3005 | -0.0712 | NA | 0.0119 | 0.0392 |
| FI | -0.0491 | NA | -0.0347 | 0.029 | -0.0187 | -0.0219 |
| FRA | -0.0432 | 0.1631 | -0.0523 | -0.0148 | 0.0137 | NA |
| HR | -0.4005 | NA | -0.2324 | -0.2321 | 0.0553 | 0.2225 |
| HU | 0.1279 | 0.3989 | -0.0292 | 0.0543 | 0.0358 | -0.0696 |
| IT | 0.1523 | 0.2041 | -0.0316 | 0.0697 | 0.053 | 0.0414 |
| LV | 0.2823 | 0.6523 | -0.0463 | 0.1409 | 0.1092 | 0.104 |
| NL | 0.0938 | 0.1723 | -0.0279 | -0.022 | 0.0381 | NA |
| NO | 0.0822 | 0.446 | -0.0107 | -0.086 | 0.0444 | 0.1056 |
| PL | 0.1323 | 0.1321 | -0.0274 | NA | 0.0563 | NA |
| SI | 0.1422 | 0.5125 | -0.0368 | 0.0816 | 0.0621 | 0.1666 |
| SK | -0.1387 | 0.4265 | -0.066 | 0.1117 | -0.0565 | -0.2172 |
| SWE | -0.083 | 0.3808 | -0.0321 | 0.0162 | -0.0309 | 0.0169 |
| SWISS | 0.1461 | -0.0304 | 0.0086 | -0.0303 | 0.0445 | NA |
| UK | 0.0055 | 0.0763 | -0.0430 | NA | 0.0140 | NA |

*S5: Mean effects of SAD, N and aggregation on species richness at small, mid and large spatial scales for the overall community (A ) as well as farmland (B) and forest ( C) guild . 'ns_bs' indicates insignificance across bootstrapping (considering only significant effects); 'NA' indicates no significance at all.*

| **A** | SAD | | | N | | | Agg | | |
|---|---|---|---|---|---|---|---|---|---|
| survey | small | mid | large | min | mid | large | min | mid | large |
| AD | 0.0311 | NA | -0.4662 | 0.1038 | NA | 0.1727 | NA | 0.2617 | ns_bs |
| AT | 0.1868 | NA | 0.5322 | 0.0253 | NA | 0.0046 | -0.4434 | -0.1414 | 0.0097 |
| BEBR | -0.0078 | NA | ns_bs | -0.0139 | NA | -0.0239 | -0.0152 | -0.0773 | -0.0268 |
| BEWA | -0.0175 | 0.2584 | 0.2754 | -0.0309 | -0.0267 | -0.0248 | 0.0474 | -0.0781 | -0.0338 |
| BG | -0.0136 | -0.4717 | -0.4951 | ns_bs | ns_bs | ns_bs | 0.1508 | 0.1901 | -0.0409 |
| CAT | 0.0772 | NA | 0.4562 | 0.0503 | NA | 0.0057 | -0.1344 | -0.1358 | 0.0268 |
| CY | 0.0176 | 0.264 | 0.4717 | 0.1172 | 0.188 | 0.1263 | -0.1223 | -0.1035 | -0.0301 |
| CZ | -0.0029 | 0.1244 | 0.1526 | 0.0207 | 0.0195 | 0.0105 | 0.0352 | -0.0806 | -0.0136 |
| DK | 0.0092 | 0.1656 | 0.206 | -0.053 | -0.0648 | -0.0473 | 0.0038 | -0.0591 | -0.0165 |
| EE | 0.0239 | 0.5443 | 0.6494 | 0.0925 | 0.1382 | 0.1092 | NA | 0.0308 | 1.00E-04 |
| ES | 0.0092 | 0.3222 | 0.3116 | -0.0814 | -0.0244 | -0.0127 | -0.042 | -0.1698 | -0.0662 |
| FI | -0.0019 | 0.0145 | -0.0606 | 0.0122 | ns_bs | ns_bs | ns_bs | -0.0397 | -0.0064 |
| FRA | -0.0123 | 0.1399 | 0.1454 | 0.0417 | 0.0054 | 0.0019 | 0.0977 | -0.0309 | -0.0623 |
| HR | 0.0689 | NA | 0.8193 | -0.2903 | NA | -0.2272 | 0.1169 | 0.0363 | 0.0165 |
| HU | 0.0829 | NA | 0.416 | -0.0396 | NA | -0.0298 | -0.2284 | -0.0663 | 0.0165 |
| IT | -0.0076 | NA | 0.1404 | 0.099 | NA | 0.02 | 0.0702 | -0.186 | -0.0186 |
| LV | 0.0144 | 0.38 | 0.4934 | 0.1021 | 0.1672 | 0.1536 | NA | -0.1454 | ns_bs |
| NL | 0.0196 | NA | 0.1738 | 0.0181 | NA | 0.0014 | 0.0712 | 0.0466 | 0.01 |
| NO | 0.0182 | 0.301 | 0.3408 | -0.0652 | -0.1199 | -0.0845 | 0.0313 | -0.1183 | 0.0026 |
| PL | 0.0945 | 0.1812 | 0.1313 | 0.076 | 0.0063 | 0.0023 | -0.0971 | 0.107 | 0.0015 |
| SI | 0.0546 | 0.4971 | 0.4763 | ns_bs | ns_bs | ns_bs | 0.0233 | -0.1004 | 0.0313 |
| SK | -0.0373 | -0.1118 | 0.1046 | ns_bs | ns_bs | ns_bs | NA | 0.2246 | ns_bs |
| SWE | 0.0522 | ns_bs | 0.3881 | -0.0125 | ns_bs | ns_bs | -0.1581 | -0.0911 | 0.0044 |
| SWISS | 3.00E-04 | NA | -0.0306 | 0.0705 | NA | 4.00E-04 | 0.0178 | 0.0153 | 0.001 |
| UK | 0.0078 | NA | 0.0577 | -0.0138 | NA | -0.0019 | 0.0097 | -0.0048 | -0.0031 |

| **B** | SAD | | | N | | | Agg | | |
|---|---|---|---|---|---|---|---|---|---|
| survey | small | mid | large | min | mid | large | min | mid | large |
| AD | 0.0022 | ns_bs | ns_bs | ns_bs | ns_bs | ns_bs | NA | -0.046 | ns_bs |
| AT | -0.0146 | 0.0308 | 0.0255 | -0.0186 | -0.0016 | -0.0011 | -0.0305 | -0.0453 | -0.0015 |
| BEBR | ns_bs | 0.0189 | -0.0061 | -0.022 | -0.046 | -0.0494 | 0.0046 | -0.0238 | -0.0116 |
| BEWA | -0.0124 | -0.0552 | -0.0509 | -0.0448 | -0.0146 | -0.0126 | 0.0301 | 0.0104 | -0.006 |
| BG | 0.0041 | -0.0785 | -0.0741 | -0.0316 | ns_bs | ns_bs | 0.0453 | 0.066 | -0.0012 |

| | | | | | | | | | |
|---|---|---|---|---|---|---|---|---|---|
| CAT | -0.0241 | -0.031 | -0.037 | -0.0194 | -0.0024 | -0.0021 | 0.0416 | 0.0187 | 0,0009 |
| CY | 0.0069 | 0.1173 | 0.1418 | 0.0296 | 0.0349 | 0.0287 | -0.0349 | -0.0768 | -0.0125 |
| CZ | -0.0049 | 0.0283 | 0.0262 | -0.0087 | -0.0038 | -0.0044 | -0.0193 | -0.0233 | -0.0038 |
| DK | 0.0028 | -0.0145 | -0.0147 | -0.0088 | -0.0052 | -0.004 | 0.0103 | -0.0092 | -0.0022 |
| EE | -0,0001 | -0.016 | -0.0169 | 0.0474 | 0.0524 | 0.0287 | NA | 0.0329 | -0,0001 |
| ES | -0.0261 | -0.003 | -0,0008 | -0.063 | -0,0009 | -0,0003 | -0.0143 | -0.003 | -0,0004 |
| FI | -0.0046 | 0.0221 | 0.0289 | -0.0108 | -0.0056 | -0.0046 | 0.0017 | -0.0209 | 0.0021 |
| FRA | -0.034 | 0.0084 | 0.0196 | -0.0287 | -0.0029 | -0.0019 | 0.0454 | -0.0408 | -0.0194 |
| HR | -0.0133 | -0.1728 | -0.1402 | -0.1377 | -0.0437 | -0.0299 | 0.0584 | -0.0416 | ns_bs |
| HU | 0.0197 | 0.0476 | 0.0463 | -0.0123 | -0.0032 | -0.0024 | -0.0293 | 0.0044 | 0.0045 |
| IT | 0.022 | 0.0985 | 0.0745 | ns_bs | 0.0028 | 0.0016 | -0.0495 | -0.056 | -0.0031 |
| LV | -0,0008 | 0.0776 | 0.1124 | ns_bs | -0.0345 | -0.0402 | NA | -0.041 | ns_bs |
| NL | -0.0357 | NA | -0.0219 | -0.0096 | NA | -0,0005 | 0.0279 | -0.001 | 0,0005 |
| NO | -0.0441 | -0.0642 | -0.054 | 0.0221 | ns_bs | ns_bs | 0.0335 | -0.0168 | -0.0031 |
| PL | -0.0083 | ns_bs | 0 | -0.0202 | -0,0002 | 0 | -0.0382 | -0.0169 | ns_bs |
| SI | 0.0155 | 0.0605 | 0.0744 | -0.0265 | -0.0042 | -0.0046 | 0.0207 | -0.0393 | -0.0077 |
| SK | 0.0041 | 0.0684 | 0.0782 | 0.0404 | 0.0374 | 0.0213 | NA | 0.0349 | ns_bs |
| SWE | 0.0089 | 0.0127 | 0.0107 | -0.0078 | -0.0037 | -0.0022 | -0.0205 | -0.0031 | 0,0006 |
| SWISS | -0.0124 | NA | -0.03 | ns_bs | NA | ns_bs | 0.0152 | 0.0098 | 0,0001 |
| UK | -0.0151 | -0.0032 | 0.003 | -0.025 | -0.0029 | -0.0018 | 0.0035 | -0.0146 | -0,0008 |

| **C** | SAD | | | N | | | Agg | | |
|---|---|---|---|---|---|---|---|---|---|
| survey | small | mid | large | small | mid | large | small | mid | large |
| AD | 0.0115 | -0.1769 | ns_bs | ns_bs | ns_bs | 0.0736 | NA | 0.0933 | ns_bs |
| AT | 0.037 | NA | 0.0489 | -0.0255 | NA | -0.001 | -0.0512 | -0.0355 | -0,0009 |
| BEBR | -0.0063 | -0.1023 | -0.1049 | -0.0141 | -0.0235 | -0.0223 | 0.0071 | -0.0353 | 0.0064 |
| BEWA | -0.0049 | 0.0167 | 0.0116 | -0.0163 | -0.0059 | -0.0023 | 0.0313 | 0.0155 | 0,0003 |
| BG | -0.0034 | -0.1079 | -0.1379 | ns_bs | ns_bs | ns_bs | 0.0062 | 0.0326 | -0.0184 |
| CAT | -0.0051 | 0.0502 | 0.0219 | 0.02 | 0.0089 | 0.0019 | 0.0194 | 0.0235 | 0 |
| CY | - | - | - | - | - | - | - | - | - |
| CZ | 0.0204 | 0.0269 | 0.0104 | 0.0339 | 0.0164 | 0.0077 | -0.0077 | 0.0175 | -0.0013 |
| DK | -0.0169 | -0.0501 | -0.035 | -0.0174 | -0.0306 | -0.0217 | 0.0364 | -0.0079 | -0.0022 |
| EE | -0.001 | 0.0361 | 0.0358 | -0.0133 | -0.0269 | -0.0167 | NA | 0.0106 | 0 |
| ES | -0.0072 | 0.0326 | 0.0308 | -0.0158 | ns_bs | -0.003 | 0.0209 | -0.0078 | -0.0032 |
| FI | 0.0156 | NA | ns_bs | -0.0145 | NA | -0.0051 | -0.0323 | 0.0204 | -0.0039 |
| FRA | 0.025 | NA | 0,0002 | 0.041 | NA | 0,0003 | 0.0411 | 0.0158 | 0.001 |

| | | | | | | | | | |
|---|---|---|---|---|---|---|---|---|---|
| HR | 0.0352 | 0.2545 | 0.2141 | ns_bs | ns_bs | 0.0665 | -0.0493 | -0.0622 | -0.0187 |
| HU | 0.0138 | 0.0424 | -0.0561 | -0.0283 | -0.0348 | -0.0278 | 0.0024 | -0.1364 | -0.0143 |
| IT | 0.0086 | 0.0153 | 0.0131 | 0.063 | 0.0328 | 0.0143 | -0.0425 | -0.0386 | -0,0003 |
| LV | 0.0185 | ns_bs | 0.0991 | ns_bs | ns_bs | -0.0382 | NA | -0.0692 | ns_bs |
| NL | -0.0219 | 0 | 0,0004 | -0.008 | -0.001 | -0,0002 | 0.0191 | 0.0086 | 0 |
| NO | -0.0189 | ns_bs | 0.1032 | -0.0336 | ns_bs | -0.026 | 0.0167 | -0.0179 | -0.0084 |
| PL | 0.0285 | 0 | ns_bs | 0.0406 | 0 | ns_bs | 0.0432 | 0.0031 | ns_bs |
| SI | 0.0115 | 0.0937 | 0.1288 | 0.0547 | 0.0574 | 0.0514 | -0.0428 | -0.0424 | 0.026 |
| SK | -0.0131 | -0.1306 | -0.1462 | -0.0647 | -0.1225 | -0.0948 | NA | 0.1461 | ns_bs |
| SWE | 0.01 | 0.017 | 0.015 | -0.0151 | -0.0109 | -0.0052 | -0.0135 | 0.0088 | -0.0015 |
| SWISS | 0.004 | NA | 0,0001 | 0.0221 | NA | 0,0001 | -0.0045 | -0.0022 | 0 |
| UK | -0.0354 | -0.0071 | -0.0015 | 0.0202 | 0.0016 | 0,0004 | 0.0356 | -0.0078 | 0 |

*S6: Mean effect of SAD, N and aggregation on species richness at small, mid and large spatial scales for farmland (left) and forest (right) birds. Color indicates the direction and strength of the effect. Gray colored countries indicate the absence of significant effect (effect size not passing Null model). dark gray colored areas were not included.*

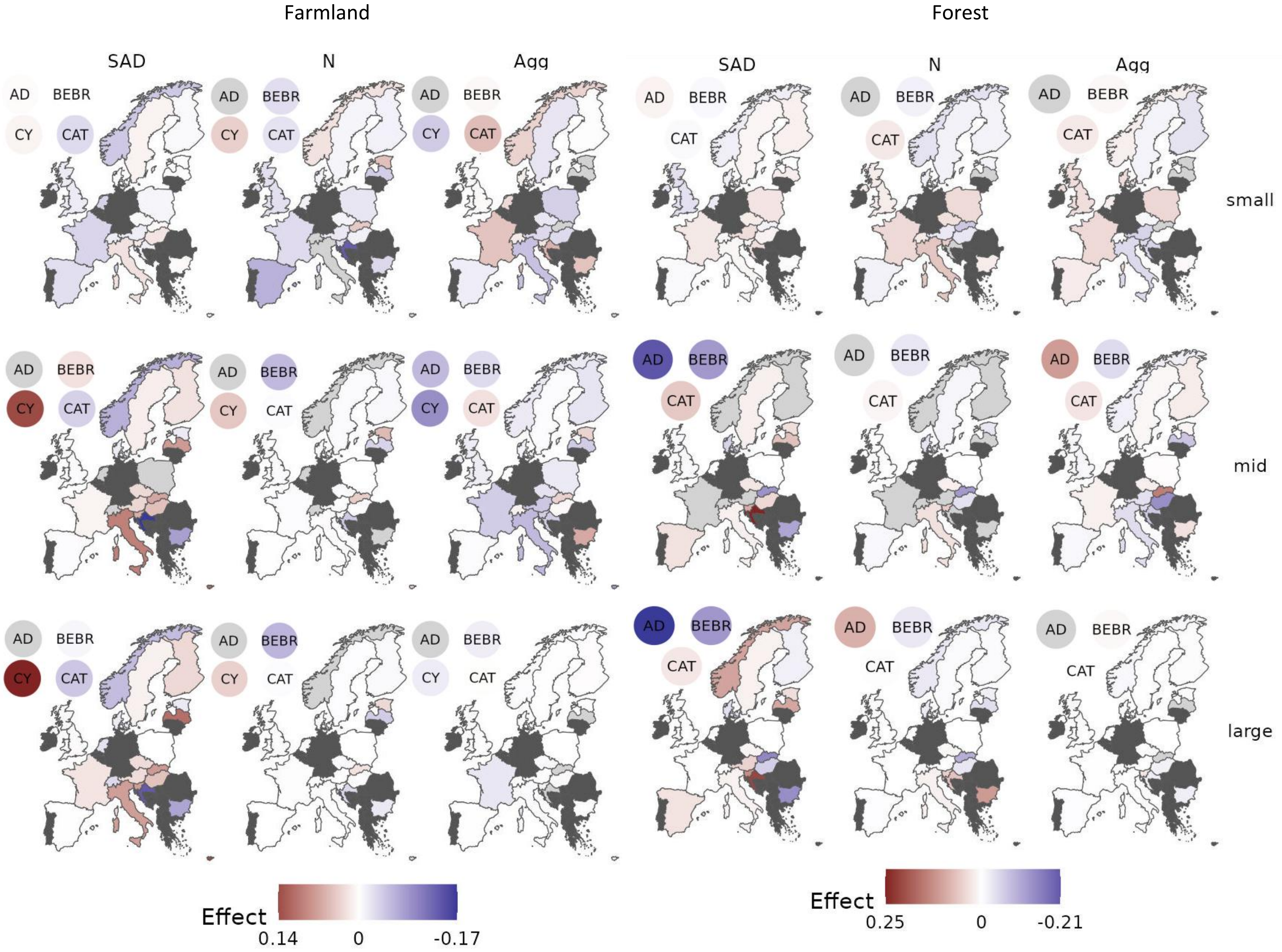

*S7: Mean effects of SAD, N and aggregation ('Agg') on species richness at small, mid and large spatial scales for within-country ecoregions. Boxplots show median and quantiles (25th, 75th), points show raw mean effects. Please note that individual scaling of x-axis to facilitate visibility. Ecoregions only appearing in one country are not shown (ALpine North (ALN), Nemoral (NEM)).*

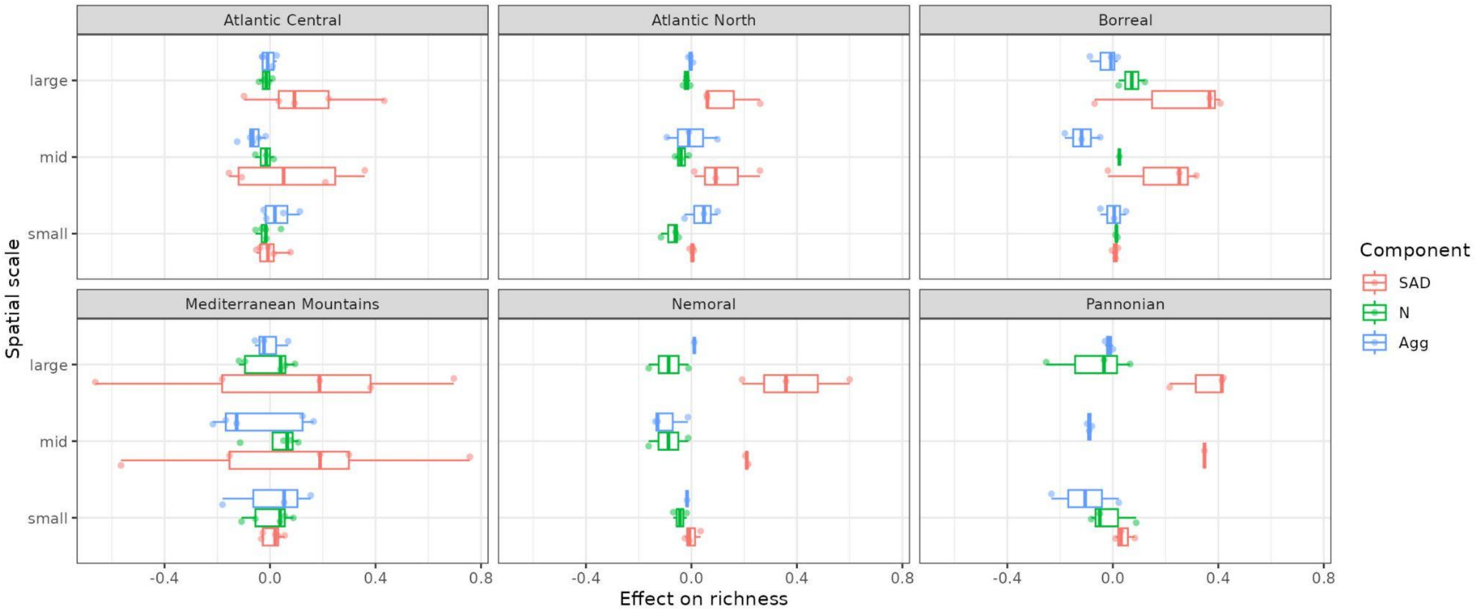

*S8: Effects of SAD, N and aggregation ('Agg') on species richness across continuous spatial scale of sampling effort as a measure of spatial scale for the overall community (top) and the farmland (middle) and forest guild (bottom). Sampling effort is measured either in number of individuals (SAD, N) or in number of sites (Agg). Color indicates the national survey. Insignificant effect values are drawn in slightly transparent color. Note that countries do not cover the same sampling effort and y-axis varies between top, mid and bottom panels for visibilit*

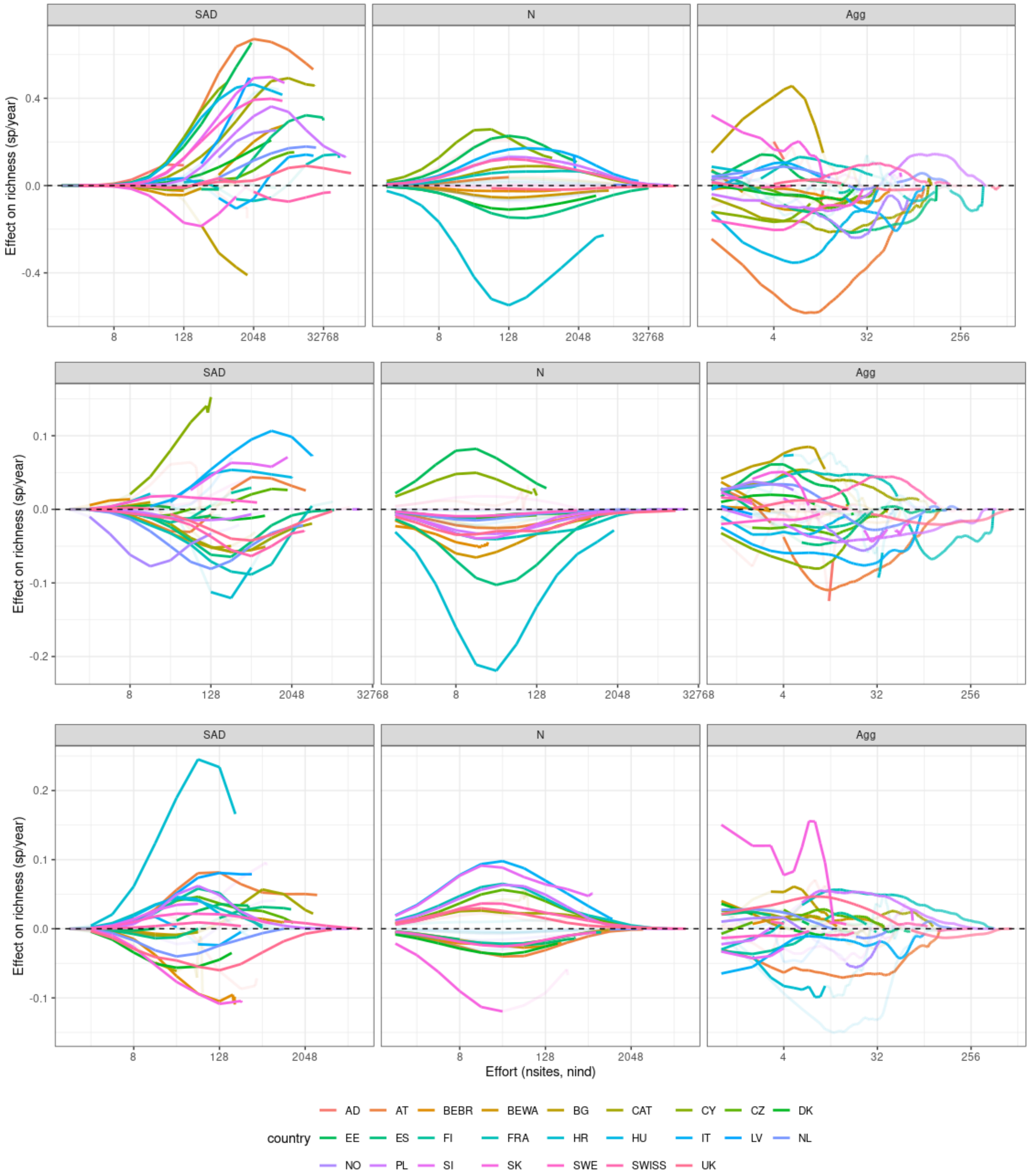
